# An Adaptive Detection for Automatic Spike Sorting Based on Mixture of Skew-t distributions

**DOI:** 10.1101/2020.06.12.147736

**Authors:** Ramin Toosi, Mohammad Ali Akhaee, Mohammad-Reza A. Dehaqani

## Abstract

Developing new techniques of simultaneous recoding using thousand electrodes, make the wide variety of spike waveforms across multiple channels. This problem causes spike loss and raise the crucial issue of spike sorting with unstable clusters. While there exist many automatic spike sorting methods, there has been a lack of studies developing robust and adaptive spike detection algorithm. Here, an adaptive procedure is introduced to improve the detection of spikes in different scenarios. This procedure includes a new algorithm which aligns the spike waveforms at the point of extremums. The other part is statistical filtering, which seeks to remove noises that mistakenly detected as true spike. To deal with non-symmetrical clusters, we proposed a new clustering algorithm based on the mixture of skew-t distributions. The proposed method could overcome the spike loss and skewed cells challenges by offering an improvement over automatic detection, alignment, and clustering of spikes. Investigating the sorted spikes, reveals that proposed adaptive algorithm improves the performance of the spike detection in both terms of precision and recall. The adaptive algorithm has been validated on different datasets and demonstrates a general solution to precise spike sorting, in vitro and in vivo.

## Introduction

Neural activity monitoring is one of the basis of understanding the brain behavior. To do so, electrophysiologists place an electrode in brain tissue to record the extracellular activity of neurons. The current flows in extracellular space would be mapped to an electric potential in the electrode. The recorded potential is a combination of multiple neurons activities corrupted by noise. A main step in analyzing the extracellular data is to differentiate among different neuron activities. In other words, we should identify each activity or spikes corresponding to each neuron. Spike sorting is the process of assigning each detected spike to the corresponding neurons. That is, spike sorting is a clustering procedure where spikes are the clustered samples.

Spike sorting algorithms usually consist of three main steps^1^. The first step is activity detection, where spikes are extracted from the bandpass filtered signal. A common way to detect spikes is to determine activities that cross a threshold^1^. The threshold is usually calculated based on the estimated noise variance, without considering dynamical changes of recorded signal, which causes spike loss or false alarms. Second, features are extracted from spike shapes. A typical method for feature extraction is principle component analysis (PCA)^2–6^. Another popular method for feature extraction is wavelet decomposition^5, 7, 8^. Finally, a clustering method would be used to assign each spike to the corresponding neuron. The clustering could be completely handled manually by human intervention, e.g. Xclust, M.A. Wilson and Offline Sorter, Plexon. In other scenarios, results are modified after an initial automated sorting^4, 9^. Modifications include merging, deleting, or splitting clusters. However, manual clustering could have more than 20% error rate^10^ and is not practical as recorded data grows up. To this end, different automatic spike sorting algorithms have been proposed in the literature. Common clustering algorithms are mixture modeling^4, 11–14^, template matching^15, 16^, and density based clustering^17^. In spite of presenting different models, their focus is more on clustering phase and supporting microelectrode array recordings with tricks like parallelism. However, the unstable clusters originated from the dynamic nature of the recorded signal is poorly investigated.

The further we go, employing high-density microelectrode arrays are getting more popular. Having multiple channels and considering the fact that the spike waveform is dependent on the distance between electrode and neuron, spike shapes of a single neuron could differ greatly. Thus, neuron shapes, and accordingly clusters, are not well formed. This problem causes more spikes to be lost and also the alignment procedure becomes more challenging. In spite of introducing different automatic sorting algorithms, the focus has been usually on the clustering phase rather than the detection. Different proposed spike sorting methods use traditional spike detection and alignment procedures and yet there exist no specific attempt for automatic and adaptive spike detection algorithm. Given the growing need for adaptive sorting methods, the concern of robust adaptive spike detection, should be addressed. To address this issues, here an adaptive detection algorithm is introduced, which consist of two main parts. The first one proposes a new alignment algorithm while the second is a statistical filtering algorithm for noise removal. The purpose of alignment is to reduce the effect of errors on determining the starting time of spike activity. In alignment, it is proposed to align detected spikes based on multiple points according to other spikes instead of just aligning according to their minimum or maximum. In addition to correcting the spike starting time, the proposed alignment, also makes the clusters more compact. Another advantage of alignment is the capability of reducing the number of principle components that are sufficient to be taken as features. For noise removal, the statistical characteristics of the spike shapes are exploited and false alarms (noises detected as spikes) are removed accordingly. The false alarm reduction is an important advantage, in that recently, we observed a great progress on timing of spiking activity. After applying the proposed adaptive detection algorithm, the spike waveforms are ready for the clustering phase.

Mixture modeling is one of the successful clustering algorithms usually used in spike sorting methods. Primary works have been focused on Gaussian mixture models, nonetheless, it has been shown that mixture of t-distribution is more powerful than Gaussian mixture in modeling neural data^14, 18^. By using mixture of Gaussian or t-distribution, we assume that the clusters are symmetrical, while as mentioned, skewed clusters are one of the challenges in the sorting algorithms. In this paper, we propose a new robust clustering algorithm based on skew-t distribution. The proposed algorithm could handle non-symmetrical clusters while preserving powerful features of symmetrical t-distribution like heavy tails.

The proposed algorithm is evaluated using synthesized and two real datasets one with and the other without ground truth information, through several experiments. Our algorithm improved the detection accuracy even in noisy channels. The proposed adaptive detection could enhance the performance of any clustering method, and is not limited to the proposed clustering algorithm. However, considering skewness made the results even more accurate. We also, developed an open-source toolbox based on the proposed method. The toolbox provides different visualizations and manual sorting functions alongside the automatic one to improve the results. Our toolbox is developed based on MATLAB and is freely available at https://github.com/ramintoosi/ROSS. Targeting false alarm and alignment, made the proposed method robust and adaptive against noise. Also, considering skewness gave us the ability to deal with not well formed cells. These challenges becomes more important when dealing with multielectrode arrays, where an adaptive detection and clustering algorithms become crucial.

## Method

In this section, we go through the details of the proposed method, which includes three main parts: preprocessing, adaptive detection, and clustering. In the preprocessing, an initial detection is performed. Next, in the adaptive detection, procedures like noise removal, alignment, and feature extraction would be handled. Finally, the spikes are clustered automatically using mixtures of skewed multivariate t distributions.

### Preprocessing

The raw data includes the spike activity combined with local field potentials (LFP), other neurons activities, and noise. Thus, the raw signal can be explained as follows:

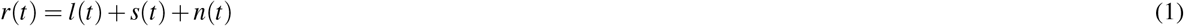

where *t* is time, *r* is the raw signal, and *l*, *s*, and *n* are the LFP, spike activity and noise, respectively. To reduce the effect of unwanted components a fourth order Butterworth bandpass filter between 300 to 3000 Hz is applied. The designed filter is a causal infinite impulse response (IIR) filter. The main advantage of IIR filters over finite impulse response (FIR) ones is their efficient hardware and software implementations, due to the fact that they reach certain specifications with a lower order. However, since the causal IIR filters have a non-linear phase response, they can alter the spike shapes dramatically or makes noises look similar as spikes^19^. This shape altering could also be the source of distraction in discriminating the pyramidal and inhibitory neuron spikes^20^ or finding relations between intra and extra-cellular activities^21^. To dispose this problem, a zero phase filter^22^ is implemented by filtering the signal in forward and reverse orders in an offline mode (for more detail please see Supplementary Appendix S1 online).

In order to detect spikes, first we need to estimate the standard deviation (std) of noise, i.e. *σ_n_*, which is calculated in the following according to Donoho’s rule^19^:

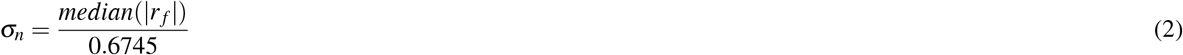

where *r_f_* is the filtered version of raw signal. The spike detection threshold is *t_s_* = 3 × *σ_n_*. It should be noted that we only use negative thresholding, i.e. detection occurs when *r_f_ < t_s_*. To increase the detection accuracy and reduce false alarms, not every threshold crossing sample would be considered as spiking activity. Given the inherent shape of an action potential, we expect the next samples pass the threshold too. Thus, a spiking activity would be detected if

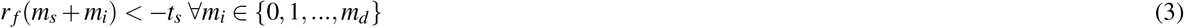

where *m_s_* is the first point that crosses the threshold and *m_d_* is the number of samples that must cross the threshold successively. Finally, the time of detected spike would be the first local minimum after *m_s_*. The local minimum is the point where there exists no point with a smaller value up to one millisecond after that. Using these local minimums reduces the early detection of spike activity caused by noise.

### Adaptive Detection

After initial detection of the spike activity, the samples of 0.5 millisecond before and one millisecond after the detected point would be captured as spike samples. In the adaptive detection phase, first it is tried to remove noise values from the detected samples which we call *statistical filtering*. Next, a novel alignment algorithm would be applied on spike samples. Finally, PCA is used adaptively for feature extraction.

#### Statistical filtering

By careful analyzing of noise samples which have been detected as spikes, we found two characteristics discriminating noise from the real spikes. The two characteristics are absolute value of average and std of samples of the waveform. For falsely detected spikes, the std of samples is large and their average is far from zero. Thus, we defined our statistical filtering based on these two parameters. To filter the false alarms, spikes whose absolute value of average and std parameters are greater than a threshold are removed. The empirically calculated thresholds are set to one and three for average and std, respectively.

#### Alignment

Now, the details of the proposed alignment algorithm would be described. Here, a general aligning algorithm, which could be used for other detection scenarios, is introduced. The proposed algorithm takes both the minimum and maximum value of the detected spike into account.

Assume that *s_pi_* is the *i*’th detected spike. First, *s_pi_* is upsampled by the factor of 10. Upsampling helped us to smoothly detect the minimum or maximum points. Then, samples are grouped based on the amplitude of their minimums or maximums, *a_min_* and *a_max_*. Samples in which *a_min_ < a_max_* are in one group and the other samples are gathered in the other one. The rest of the algorithm would be applied on each group separately. Assuming the group where *a_min_ < a_max_*, the histogram of the indices of the maximum values would be calculated as shown in Fig. 1(b). The histogram is smoothed using a cubic spline interpolation. Then, the *n_peak_* greatest peaks in the histogram would be considered as aligning points (1(b)). For each sample in the group, if the maximum of the sample could reach the nearest aligning point by the maximum shift of *n_shift_* point, it would be shifted to be aligned with the nearest aligning point (1(c)). The aligning algorithm is illustrated in Fig. 1(a).

**Figure 1.**
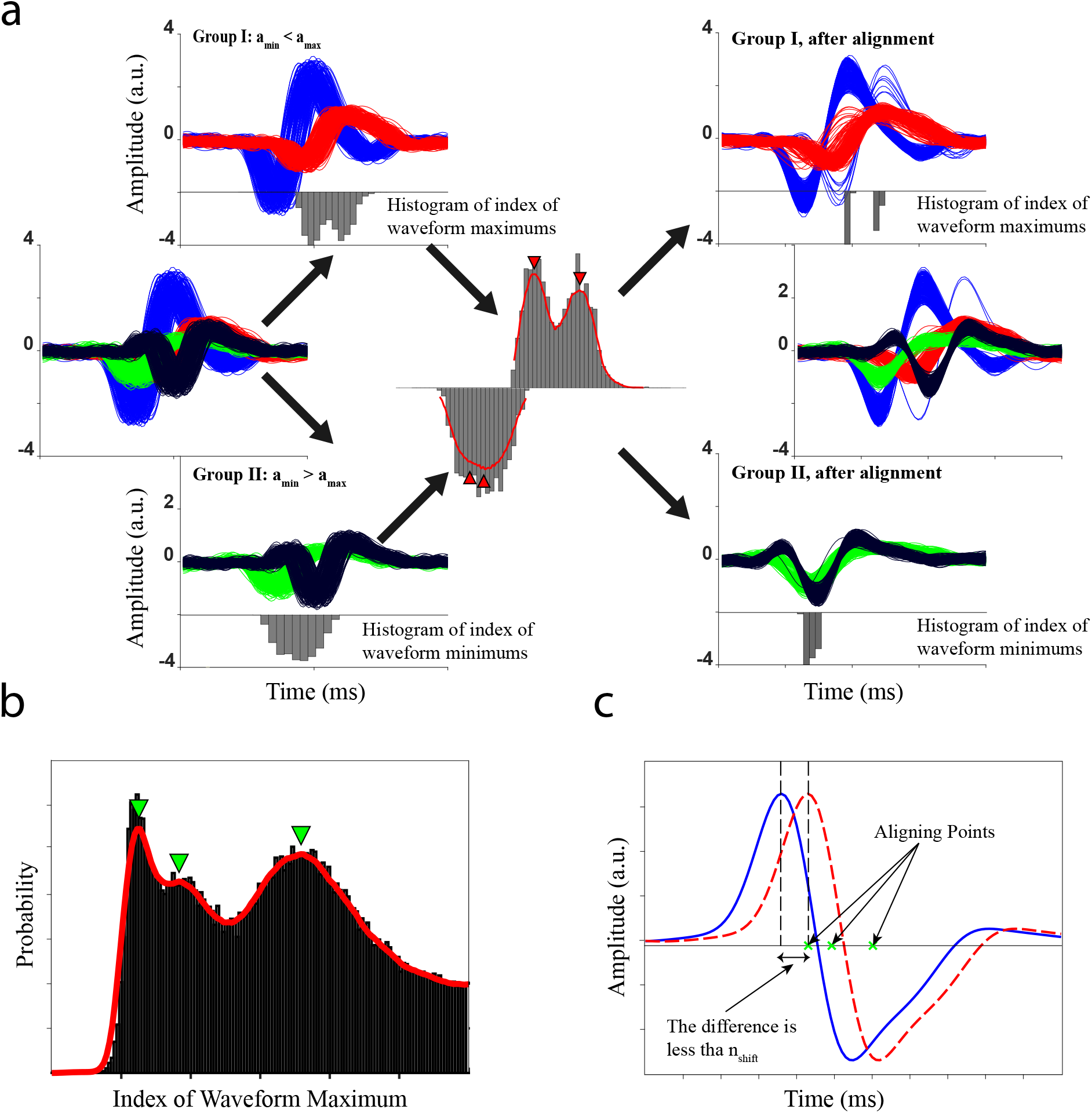
An intuitive example of the proposed alignment algorithm. (a) Alignment of synthesized waveforms. The synthesized waveforms are designed to form four distinct shapes. The waveforms are grouped according to the amplitude of their maximums and minimums. Then, the peak of the smoothed histogram of extremum indices is found, as the aligning points. Finally, the waveforms are aligned regarding the aligning points. (b) The histogram of peak locations in waveforms. The smoothed cubic spline fitted to the histogram is shown by the red line. The locations of three local maximums are indicated using green arrows. (c) The blue solid line is the original waveform. The aligning points, corresponding to the local maximums in (b) are also depicted using green crosses. If the difference between the peak location of the waveform and the nearest aligning point is less than *n_shift_*, the waveforms is aligned by an appropriate shift, forming the aligned waveform illustrated by dashed red line.

There exist two parameters in the alignment process, i.e. *n_peak_* and *n_shift_*. Here, we discussed about the extreme cases of these two parameters. When, *n_peak_* is one, the algorithm seeks to align all spikes into one unique point. In this way, if the difference between two neuron spikes is the index of their extremum, then the algorithm misleads the clustering process by removing a discrimination characteristic. The other extreme case is when *n_peak_* is equal to the number of samples. In this scenario, the algorithm does not align any waveform. Thus, increasing the value of *n_peaks_*, reduces the aligning effect. The other parameter is the maximum allowed shift for aligning a sample, i.e. *n_shift_*. Small values of this parameter reduce the effect of the algorithm. Also, large shifts dramatically change the spike shapes which is not desirable. After alignment, the tails of the spike samples, which does not carry important information about the spike shape would be eliminated.

#### Feature extraction

Applying PCA, the spike samples are projected into a low dimensional space for the feature extraction process. Instead of using a fixed number of components, an adaptive method is used. Assume *λ_i_* is the *i*’th greatest principle component variance. We select *n_c_* components where *n_c_* is the smallest solution that satisfies the following condition:

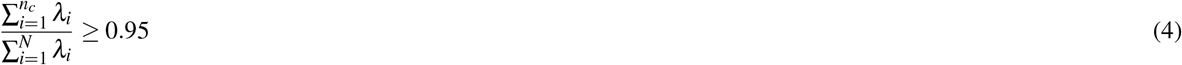

where *N* is the total number of components. Intuitively, the new low dimensional space preserves 95% of the variance in the original high dimensional space to describe spike shapes. This method allowed the algorithm to adaptively choose the dimension of the feature space base on the spike shape variations. However, choosing a large number of components may affect the clustering process because of the *curse of dimensionality*. Therefore, if the number of selected components by (4) exceeds a predefined value, *N_pca_*_−*max*_, the algorithm chooses the first *N_pca_*_−*max*_. Throughout this paper, we fixed *N_pca_*_−*max*_ = 15.

### Automatic Spike Soring Using Mixture of skew-t distribution

In this section, we propose a new automated spike sorting algorithm based on mixtures of skew-t distribution. The expectation maximization (EM) algorithm applied in this section is based on the work of Cabral *et. al.*^23^. In their work, the skew normal independent (SNI) family is discussed. The proposed algorithm is based on the skew-t distribution which is a member of this family. Here, we only discuss about the results of their work for the skew-t distribution.

To define the multivariate skew-t (ST) random variable statistics, first we need to define the multivariate skew normal (SN) one. A p-dimensional random variable X follows the SN distribution, if its distribution is given by:

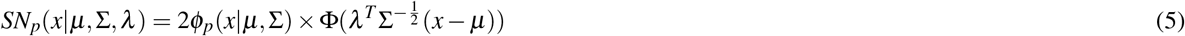

where *μ* is the *p* × 1 location vector, Σ is *p × p* positive definite dispersion matrix, and *λ* is the *p* × 1 skewness parameter vector. *ϕ_p_*(. |*μ,* Σ) is the probability density function (PDF) of the p-dimensional normal random variable, and Φ(.) stands for the standard univariate normal cumulative distribution function (CDF).

Assume the following random variable, *Y*,

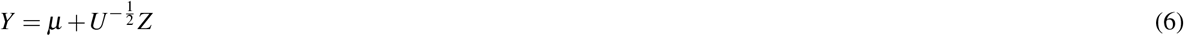

where *μ* is a p-dimensional location vector, *Z ~ SN*(0, Σ*, λ*), and *U* is a positive random variable, independent of *Z*, with PDF h(.|v). If *U* follows a *Gamma* 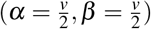 distribution, i.e.

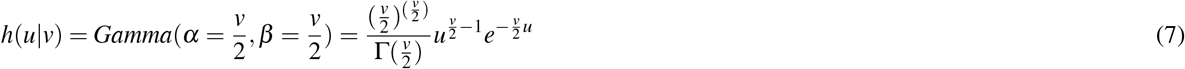

where Γ(.) is the Gamma function, then, *Y* follows a multivariate ST distribution with the location vector *μ*, dispersion matrix Σ, skewness vector *λ*, and *v* degrees of freedom. The distribution of *Y*, could be simplified based on (5) – (7) as follows:

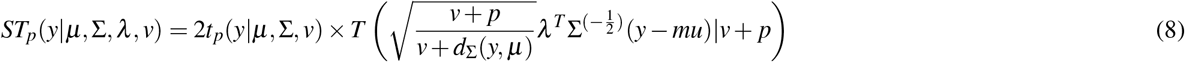

where *t_p_*(.|*μ,* Σ*, v*) denotes the pdf of the p-dimensional student-t distribution with the mean *μ*, dispersion matrix Σ, and *v* degrees of freedom. Also, *T* (.|*v* + *p*) is the standard univariate student-t CDF with *v* + *p* degrees of freedom. *d*_Σ_(*y, μ*) = (*y* − *μ*)^*T*^ Σ^−1^(*y* − *μ*) is the Mahalanobis distance between *y* and *μ*.

#### Mixture of multivariate skew-t distribution

In mixture models, we assume that each spike sample, denoted as *s_i_*, originates from a finite set of known distributions (or components) with unknown parameters. First, it is assumed that the number of distributions, *g*, is known. In the next subsection, when mixture model is used for clustering, *g* would be determined automatically. Assuming that the spikes are *i.i.d* random variables denoted by *S*, the likelihood function can be written as:

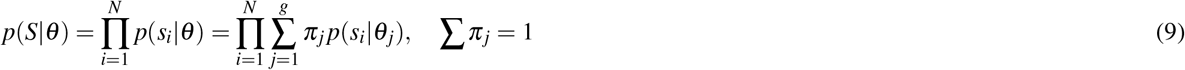

where, *θ_j_* is the distribution parameter set, and *π_j_* is the mixing probability. In the proposed method, components are ST distributions (i.e. *p* = *ST_p_*) with the parameter set *θ_j_* = {*μ _j_,* Σ _*j*_, λ_j_, *v*}. To find the optimum set of parameters, we employ EM algorithm. The details of the EM formulation is provided in Supplementary Appendix S2 online.

To find the optimum parameters, the algorithm iteratively does the expectation and maximization steps respectively, until a stop criteria is met. Intuitively, the algorithm should stop when the parameters reach to a stable value in the updating procedure. Mathematically, the algorithm stops, if the following condition is met:

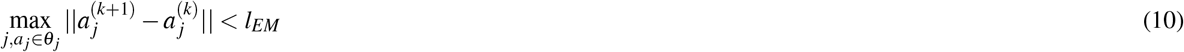

The condition captures the maximum change among all parameters of all components, and then checks if it is smaller than a predefined threshold, i.e. *l_EM_*. The EM-algorithm is summarized in Algorithm 1, where *θ_init_* is the initial values of the parameters.

#### Clustering method

In the previous section, we used the EM algorithm to find the optimum parameters of the components. In the EM-algorithm, it is assumed that the number of components, i.e. *g* is known. But in the automatic spike sorting problem, the number of components or neurons is unknown and the algorithm have to determine it. The EM-algorithm would be referred to as *θ_out_* = *STEM*(*θ_init_, g*), where *θ_init_* and *θ_out_* are the initial and output (optimum) parameters as shown in Algorithm 1. Suppose that the clustering algorithm searches the best value of *g* within the interval [*g_min_, g_max_*]. A simple way is to run STEM for *g* = *g_min_,…, g_max_* independently. Besides, the random initialization of parameters could be used for each run. Then, a criterion determines the optimum value of *g*. Here, to speed up the algorithm, we exploit the output of the previous run of the STEM to initialize the parameters of the next run.

##### Algorithm 1

EM-algorithm for mixture of skew-t distributions (STEM)

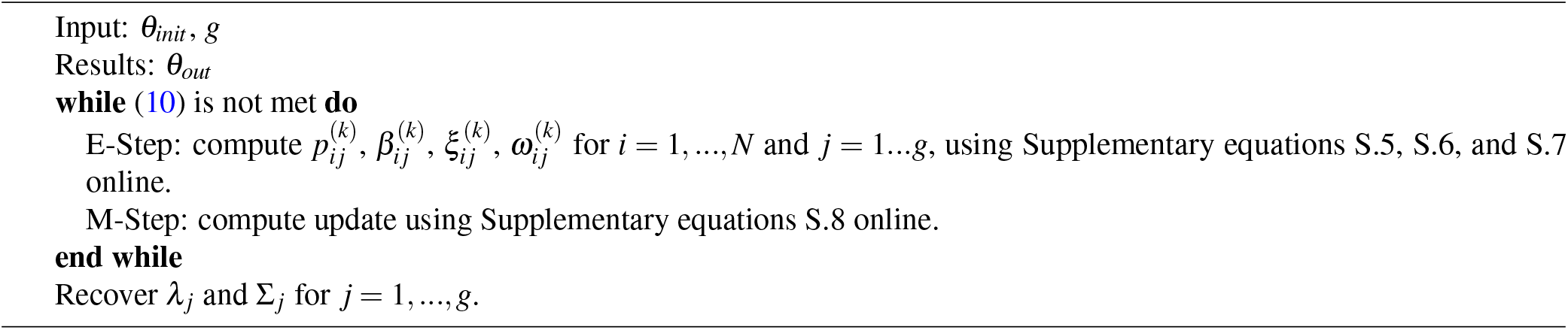

The proposed clustering algorithm starts from *g* = *g_max_* and continues the search toward *g* = *g_min_*. In the first run, when *g* = *g_max_*, we use the fuzzy c-means clustering (FCM) algorithm to calculate *θ* instead of using random initialization. The FCM algorithm outputs the cluster centers and a fuzzy partition matrix, *U*, where *U_i j_* is the probability that the *i*’th spike belongs to the *j*’th component. *μ* would be initialized by centers of the FCM results. The dispersion matrix is

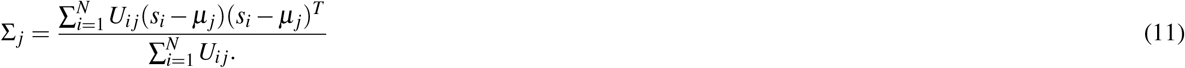

Also, the initial value of the skewness vector, *λ* is,

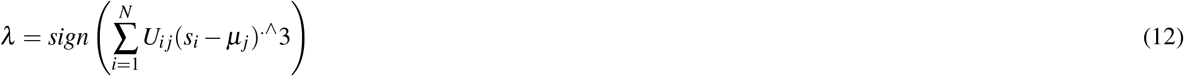

where .^ is the element wise power.

After running the STEM algorithm, the estimated parameters would be exploited to initialize the next run. Suppose that *θ_out,k_* is the estimated mixture parameters in the *k*’th step. The initial values for the next run would be calculated as follows:

- Find the component with minimum mixing probability, *m* = arg min_*l*_ π_*l*_
- Remove *π_m_* and normalize the remaining mixing probabilities such that, ∑_*l*_ π_*l*_ = 1.
- Remove *μ_m_*, Σ_*m*_, and *λ_m_*.
- Use the output degree of freedom, *v*, to initialize the next one.
- Set *g* ← *g* − 1.

In order to find the best value of number of components, *g*, we use the likelihood function in Supplementary equation S.3 Online. Assume *L_g_* denotes *ℓ_c_*(*θ* |*s, t, u, z*) when the number of components is *g*, then the optimum value of *g*, i.e. *g_opt_* is,

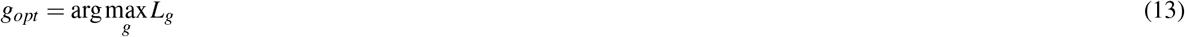

Also, One can terminate the algorithm if increasing of *L_g_* is observed. Finally, the cluster index of each spike would be obtained as follows:

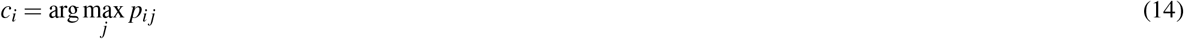

where *c_i_* is the cluster index of *s_i_*. The proposed clustering algorithm is summarized in Algorithm 2.

## Results

Here, the performance of the proposed method is investigated using three different datasets, including one simulated and two real datasetss. First, the details of the datasets are described. Then, we go through the details of the experiments and the results are discussed. Through this section, the proposed method is compared with the method proposed by Shoham^14^. From now on, the method of Shoham^14^ and the proposed method are referred to as MTD (mixture of t-distributions), and MSTD (mixture of skew-t distributions). In the following experiments, all statistical comparisons are calculated using signed-rank test.

### Algorithm 2

Clustering Algorithm

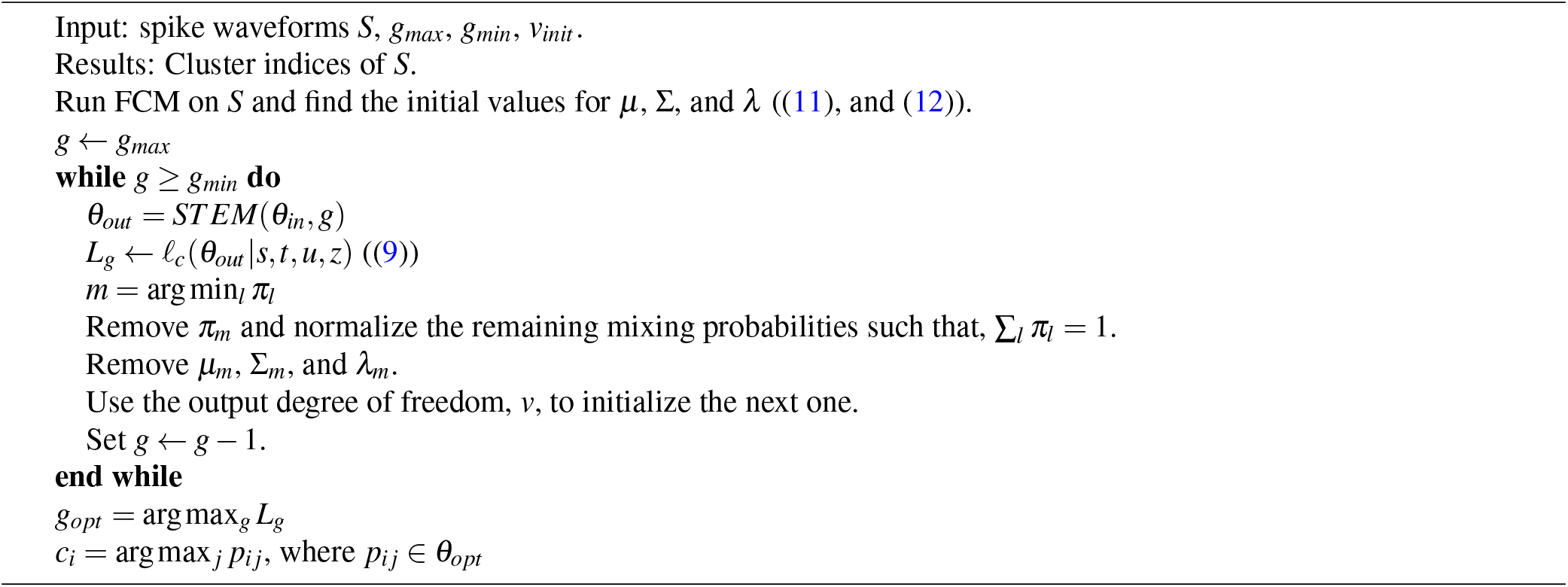

### Datasets

A synthetic dataset is used to show the correctness and efficiency of the proposed method. To simulate neural recording data, the tool noisy spike generator provided by Smith and Mtetwa^24^ is employed^25^. Using this tool, 200 sessions of recording are simulated. Each session contains 200 seconds of recording with sampling rate of 40KHz. Four neurons with Poisson distribution in each session with refractory period of one millisecond is set. The other options are leaved as their defaults. This resulted in approximately 30000 spikes in each session.

For the first real dataset, we employed the simultaneous intracellular and extracellular recordings from hippocampus region CA1 of anesthetized rats, provided by Henze *et al*.^21, 26^. Sessions with appropriate intracellular recording were selected manually. Here, the intracellular recording is exploited as the ground truth to evaluate the performance of the proposed spike detection and preprocessing. The third dataset is a recording of a macaque. These sessions are collected from inferotemporal cortex area, while monkey does a rapid serial visual presentation task. It consists of 200 sessions. The sampling rate is 40KHz.

### MSTD Methodology Overview

Fig. 2, graphically illustrates the procedure of the proposed method. In the preprocessing step, first, signal is bandpass filtered using zero phase filtering method. Next, using a simple thresholding, an initial detection of spikes is performed. Threshold is calculated as three times the standard deviation of noise. At this point, a bunch of detected waveforms is ready for the proposed adaptive detection algorithm. Here, statistical filtering tries to reduce the system false alarm by removing noises that are mistakenly detected as spike. For this sake, the waveforms that their average or standard deviation exceeds a predefined value would be removed from data. Then, the proposed alignment algorithm seeks to align the waveforms based on the histogram of the extremums. The main point of our alignment algorithm is to use different points to align a waveforms instead of only considering its minimum or maximum. The last step in adaptive detection is the feature extraction, where we adaptively choose a number of principle components as feature. Finally, the samples are fed to the proposed clustering method based on the mixture of skew-t distributions. The proposed clustering method could handle skewed cells and automatically finds the best number of neurons.

**Figure 2.**
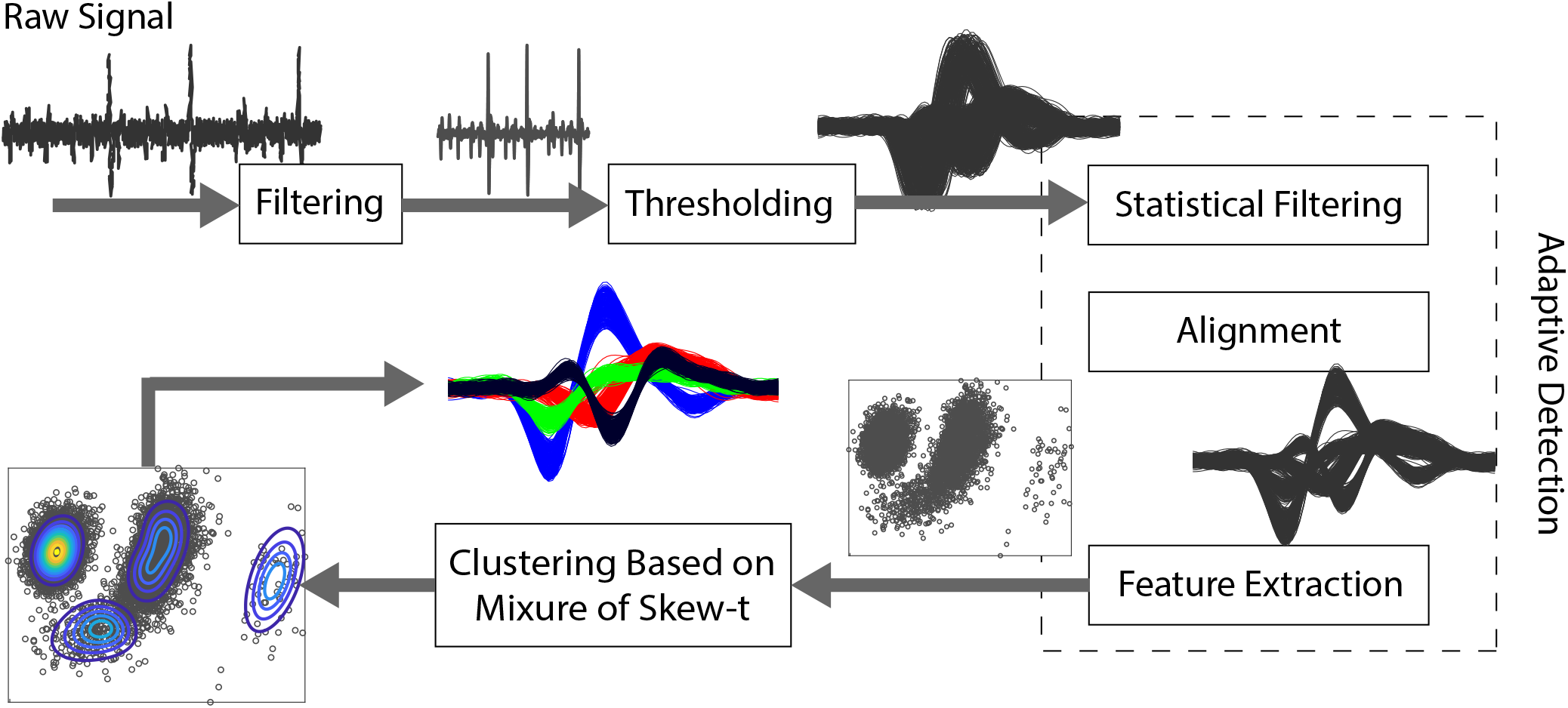
Illustration of the procedure of the proposed automatic sorting method. In the proposed method, first the signal is filtered. Then, an initial detection is performed applying a simple thresholding method. Thereafter, the proposed adaptive detection algorithm takes the initially detected spikes and performs a set of algorithms to improve the detection results and make them ready for the clustering phase. In adaptive detection, first the noise which is detected as waveforms are removed using proposed statistical filtering. Then the proposed alignment algorithm, seeks to align the waveforms based on the histogram of the extremums. Then, an adaptive number of principle components are considered as feature. Finally, the samples are fed to the clustering method based on the mixture of skew-t distributions.

### Application and Validation of Adaptive Detection on Synthesized Dataset

In the first experiment, we used the simulated dataset to evaluate the proposed detection and preprocessing algorithms. The true spikes of a sample session are visualized using the first two components of PCA in Fig. 3(a). As can be seen, there exist four distinct groups of spikes, as illustrated in Fig. 3(a)(top) with different colors. The distribution of spikes is also illustrated in Fig. 3(a)(bottom). For better evaluation, the population of each group is chosen from a range of highly populated to the sparse one. In this experiment, eight Gaussian noise signals with different powers is added to each recording to produce noisy signals with controlled signal to noise ratios (SNR). Then each signal is filtered and spikes are detected as described in the detection algorithm. A sample of clean, noisy and filtered signals are illustrated in Fig. 3(b). Each noisy recording, in each SNR is fed to the detection process of MTD and MSTD. A detected spike is considered as a true positive if there exist at least one spike in its one millisecond vicinity in the ground truth. To evaluate the performance, the *precision* and *recall* metrics are calculated. Precision indicated how many of the detected spikes are actually a true spike and could be calculated using the following formula:

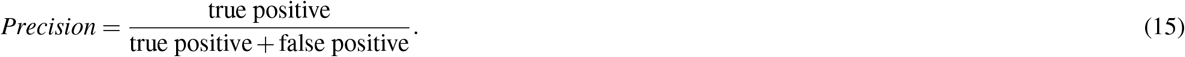

**Figure 3.**
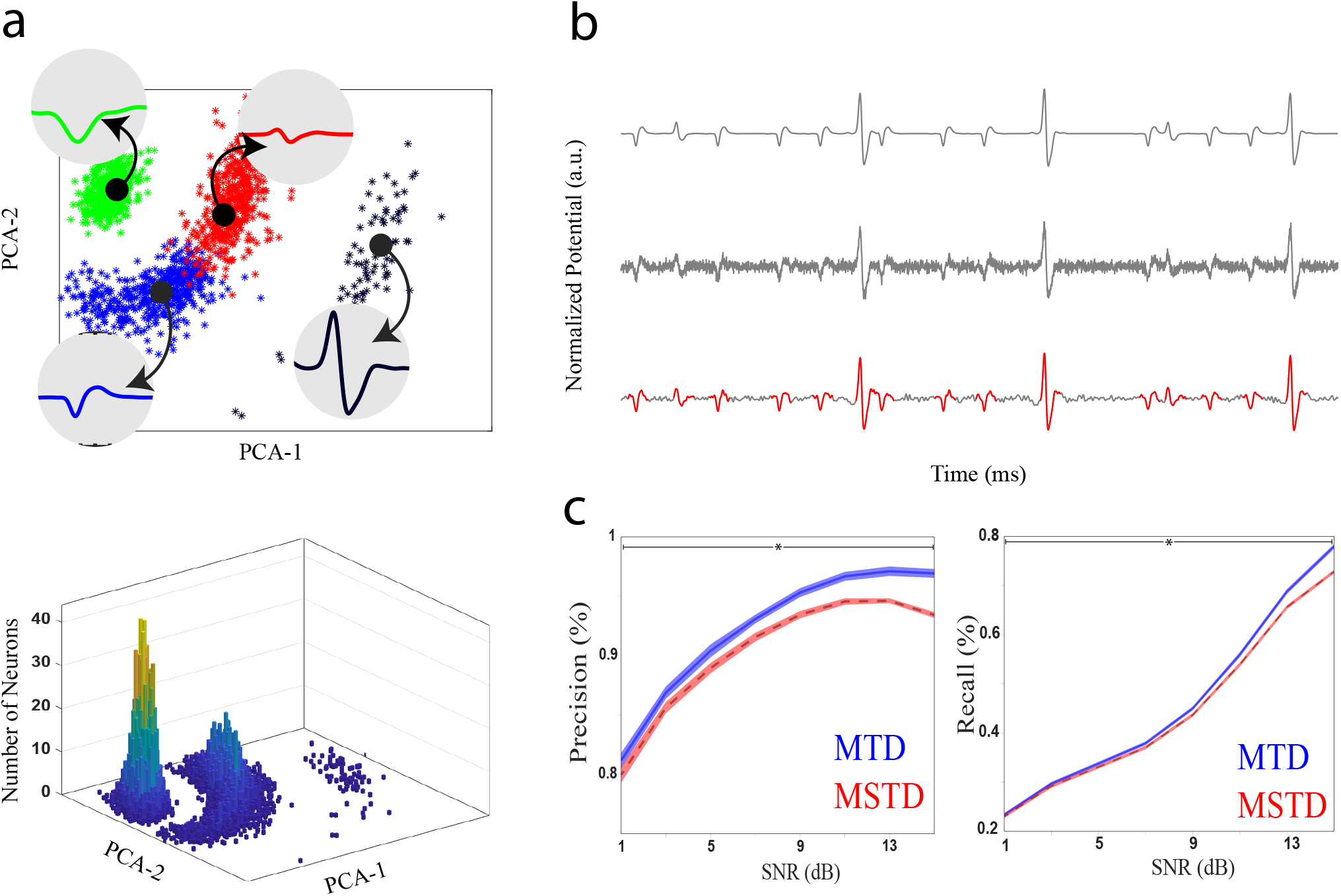
Evaluating the MSTD method using simulated dataset^24, 25^ in the detection phase. The dataset contains 200 sessions of 200 second recording with sampling rate of 40KHz. (a) A sample session is visualized using its two first principle components (top). There exist four distinct neurons distinguished by four colors. The average waveform of each neuron is also indicated in 1.2 millisecond interval. A sample distribution of spike waveforms is also illustrated (bottom). The x and y axes are the first two principle components of waveforms and the z axis is the relative frequency. (c) A sample of clean recording data (top), noisy data (middle), and filtered data (bottom). (d) Evaluating the proposed detection, alignment and statistical filtering algorithm. The precision (left) and recall (right) of MSTD in the detection phase is compared with MTD. The shaded area is the standard deviation. The horizontal line with star shows where the values differ significantly.

Recall shows how many of the true spikes are detected by the algorithm. The recall can be stated as follows:

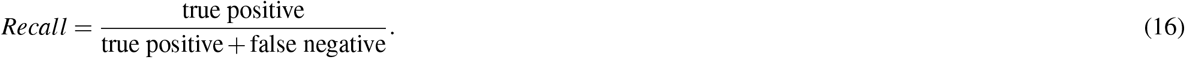

The high precision means less false alarms and high recall means less missed spikes. The average precision and recall of MTD and MSTD methods for each SNRs are illustrated in Fig. 3(c). The shaded area shows the standard deviation of each point. As this figure illustrates, the MSTD method improves the performance of the detection phase in terms of both precision and recall in high SNRs. In low SNRs, however, only precision shows an improvement over MTD method. In SNR= 15 dB, MSTD increases the precision and recall of MTD by 6% ± 0.1% and 7% ± 0.1%, respectively. This confirms the effectiveness of our adaptive detection algorithm.

### Application and Validation of proposed clustering algorithm on Synthesized Dataset

After the detection phase, the detected spikes are fed to the clustering algorithms of MTD and MSTD. For a fair comparison, both methods are applied on the same set of detected spikes in each session with the same SNR. One example session with fitted distribution applying MSTD is illustrated in 4(a). The average number of automatically found clusters by each method in different SNRs alongside their standard deviations is demonstrated in Fig. 4(b). As can be seen, in a large SNR interval (5dB to 13dB) the average number of clusters is close to the real number of clusters (each recording has four neurons) with less standard deviation. Thus, MSTD determines the number of clusters better than MTD. To evaluate the clustering quality and also considering the ground truth, the accuracy and purity metrics are employed. For accuracy, the label of each cluster is determined by the dominant neuron in that cluster. The dominant neuron in each cluster is the neuron that has the most number of spikes in that cluster. For purity, the percent of the number of dominant neuron samples in a cluster is considered as the purity of a cluster. The average purity of all clusters is calculated as the clustering purity. The average accuracy and purity is demonstrated in Fig. 4(c-d). As shown, in SNRs greater than five, MSTD outperforms MTD in terms of both accuracy and purity. At SNR 11, the accuracy and purity of MSTD is 4% ± 3% and 7% ± 4% greater than that of MTD, respectively. As a consequence, the MSTD algorithm better clusters the detected spikes.

### Application and Validation of MSTD on Real Dataset with Ground Truth

In the next experiment, the MSTD and MTD methods are examined using a real dataset with ground truth for the time of spikes (rat hippocampus). The extracellular recordings are fed to the detection algorithms of MTD and MSTD methods. Here, a precision-recall plot is employed to perform a comparison between two methods for a range of precision and recall values, simultaneously. In this way, two methods could be compared in low and high precision or recall states. To achieve this, the threshold is selected as a variable coefficient of the estimated noise power, i.e. *t_s_* = *c_t_ × σ_n_* where *c_t_* ∈ {1, 1.5, 2, 2.5, 3, 3.5, 4, 4.5, 5}. The result is depicted in Fig. 5(a). The standard deviation for each average value of precision and recall is also indicated using horizontal and vertical error bars, respectively. As can be seen, MSTD reaches a higher precision for the same value of recall and vice versa. This improvement is the result of our detection and preprocessing algorithms. The reason of the low precision values in 5(a) is illustrated in Fig. 5(b). As indicated, the ground truth achieved by intracellular recording only contains the spikes of one neuron; however, in a recording there may exist several neurons which explains the low values for precision. In this example, our manual sorting showed three kinds of neurons in this recording.

**Figure 4.**
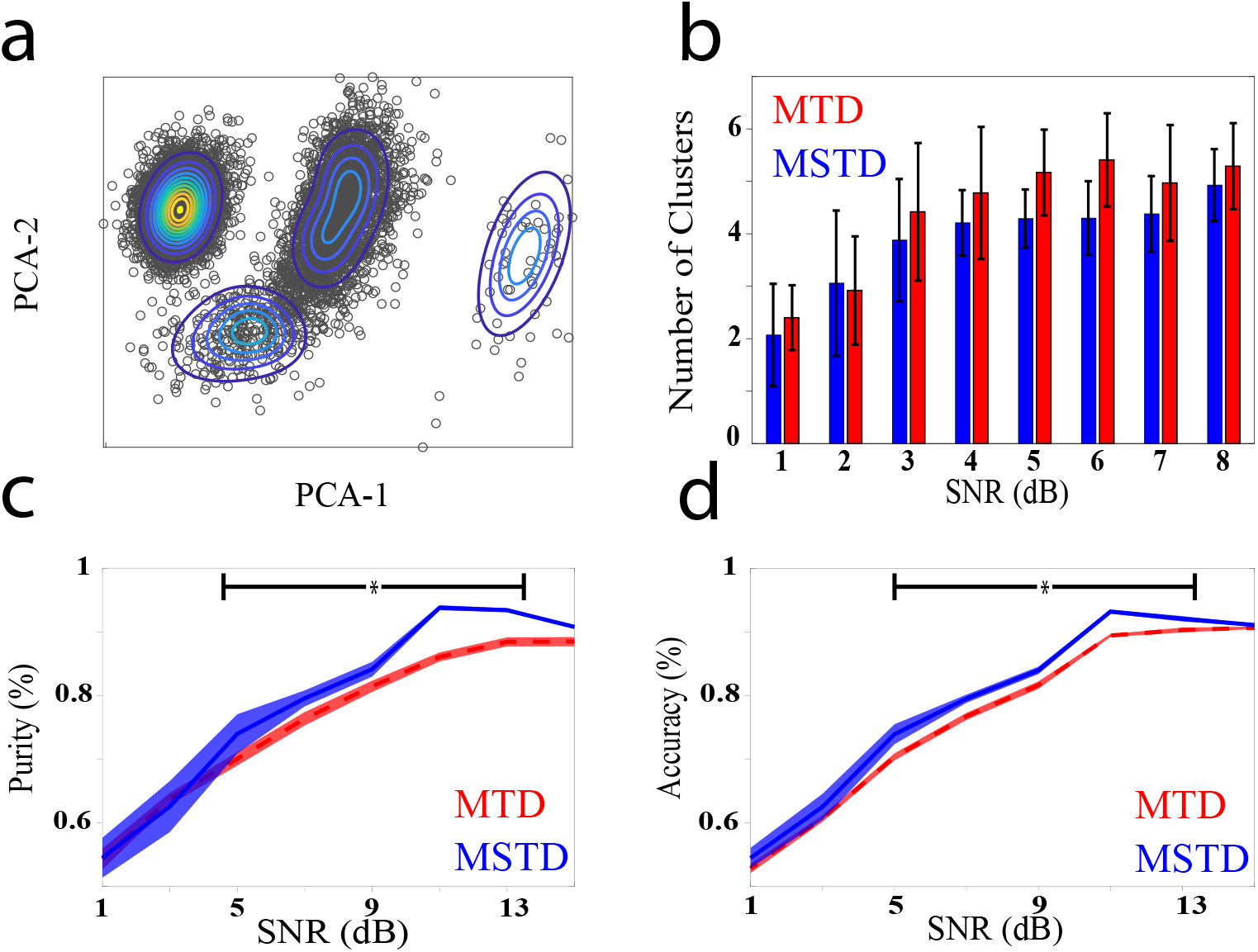
Evaluation of MSTD method using simulated dataset^24, 25^ in clustering phase. (a) An example of MSTD fitted on a simulated session. (b) The average number of clusters determined by each method for different SNRs. The standard deviation is shown as error bars. (c-d) The accuracy and purity of clustering outcomes for MSTD and MTD methods. The shaded area is the standard deviation. The horizontal line with star shows where the values differ significantly.

**Figure 5.**
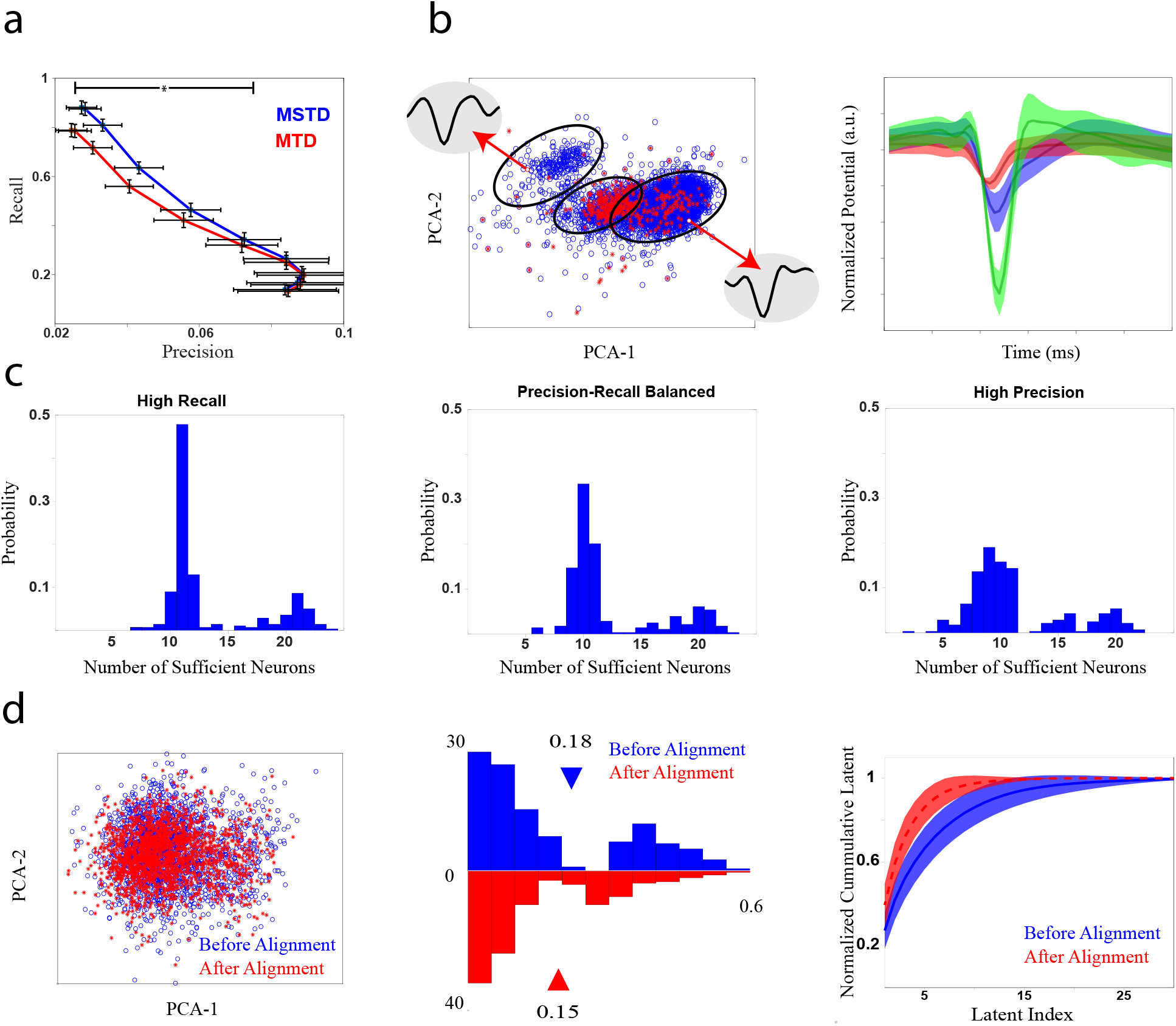
Comparison of MTD and MSTD using real dataset of simultaneous intracellular and extracellular recordings of rat hippocampus^26, 27^. (a) Precision-recall curve of detection results. To achieve this curve the detection threshold is varied proportional to the noise power. The standard deviation of the precision and recall is indicated using horizontal and vertical error bars, respectively. As can be seen, the precision values are small. The horizontal line with star shows where the values of both recall and precision differ significantly. (b) It shows the reason of small values of precision. A recording sample is visualized using the two first principle components (left). The blue circles show the detected samples, and the red stars show the spikes detected using the ground truth. In the right, the average waveforms of neurons of this sample is illustrated. The clusters are formed using simple manual clustering. The shaded area states the standard deviation. (c) The histogram of the number of principle components is needed to describe the spike waveforms, *n_c_*, calculated by equation (4), for three situations: high recall (left), which is the point with most recall value in (a), precision-recall balanced (middle), which is the point in the middle of high recall and high precision points in (a), and high precision (right), which is the point with the most precision value in (a). (d) The effect of alignment on the cluster compactness. A sample of ground truth spikes of two recordings are visualized using the first two principle components (left). The blue circles and red stars show data before and after the proposed alignment, respectively. The histogram of within distance of ground truth spikes before and after applying the proposed alignment (middle). The left side of equation (4) is shown before and after alignment (right). The shaded area is the standard deviation.

In the feature selection, the condition to select a variable number of features, *n_c_* (see equation (4)) is introduced. Here, using a real dataset, we showed the reason of choosing a variable number of featues. The distribution of the number of components that satisfies (4), i.e. *n_c_*is depicted in Fig. 5(c). The results are shown for three different situations: i) high recall (*c_t_*= 1), ii) precision-recall balanced (*c_t_*= 3), and iii) high precision (*c_t_*= 4.5). As seen, in all situations *n_c_*varies in a wide range, demonstrating the variability of spike waveform changes from one recording to another; thus, an adaptive method to select the number of components is required.

Next, we examined the effect of the alignment algorithm on the spike waveforms. The spike waveforms are detected using the provided ground truth times; then two kinds of alignment algorithm are applied. The first one is to align waveform according to their minimums, and the second one is the proposed algorithm. Two samples of neuron spikes are indicated in the left side of Fig. 5(d). The alignment algorithm seemed to make the neuron spikes more compact. To evaluate this, the distribution of within distance of clusters are depicted in the middle plot of Fig. 5(d). The alignment algorithm significantly (*p* = 5×10^−27^) reduced the within distance of the clusters, which means it makes the waveforms of one neuron more similar. Nevertheless, since the intracellular recording provided the ground truth for one neuron, we cannot examine the effect of alignment on between distances of neurons. Another way to examine this effect is the contribution of each principle component in the total variance. The right side of Fig. 5(d) shows the left side of equation (4) for the two alignment algorithms. This figure shows that the proposed algorithm makes the waveforms to concentrate their variances in the lower dimensions, which means that spikes could be better described in lower dimensions.

### Application and Validation of MSTD on Real Dataset without Ground Truth

In the final experiment, we examined the MSTD algorithm with our dataset which had no ground truth. To evaluate the effect of the introduced statistical filtering algorithm, three recording samples are shown in Fig. 6(a). The red stars are the detected noises using statistical filtering. Samples of waveforms detected as noise are also indicated. In the PCA space, the detected waveforms are sparsely located away from the concentrated location of spikes. Their waveforms are also highly different from other spikes stating the effectiveness of the proposed statistical filtering algorithm. Since, there exists no ground truth for this dataset, we used three metrics to evaluate the clustering quality: i) sum of square errors (SSE), ii) Calinski-Harabasz criterion^28^, and iii) Davies-Bouldin criterion^29^. As shown in Fig. 6(b), the clustering quality of MSTD is significantly better than MTD in terms of SSE (*p* = 5 × 10^−25^) and is significantly worse than MTD in terms of Calinski-Harabasz criterion (*p* = 10^−6^). Also, there exists no significant difference between two methods from the Davies-Bouldin criterion point of view (*p* = 0.44). Thus, these blind criteria could not actually show a significant superiority simultaneously. The distribution of the number of clusters determined by both algorithms is plotted in Fig. 6(c). As can be inferred, MSTD tends to have more clusters in this dataset. Also, MTD results in only one cluster in about twenty percent of the recordings.

**Figure 6.**
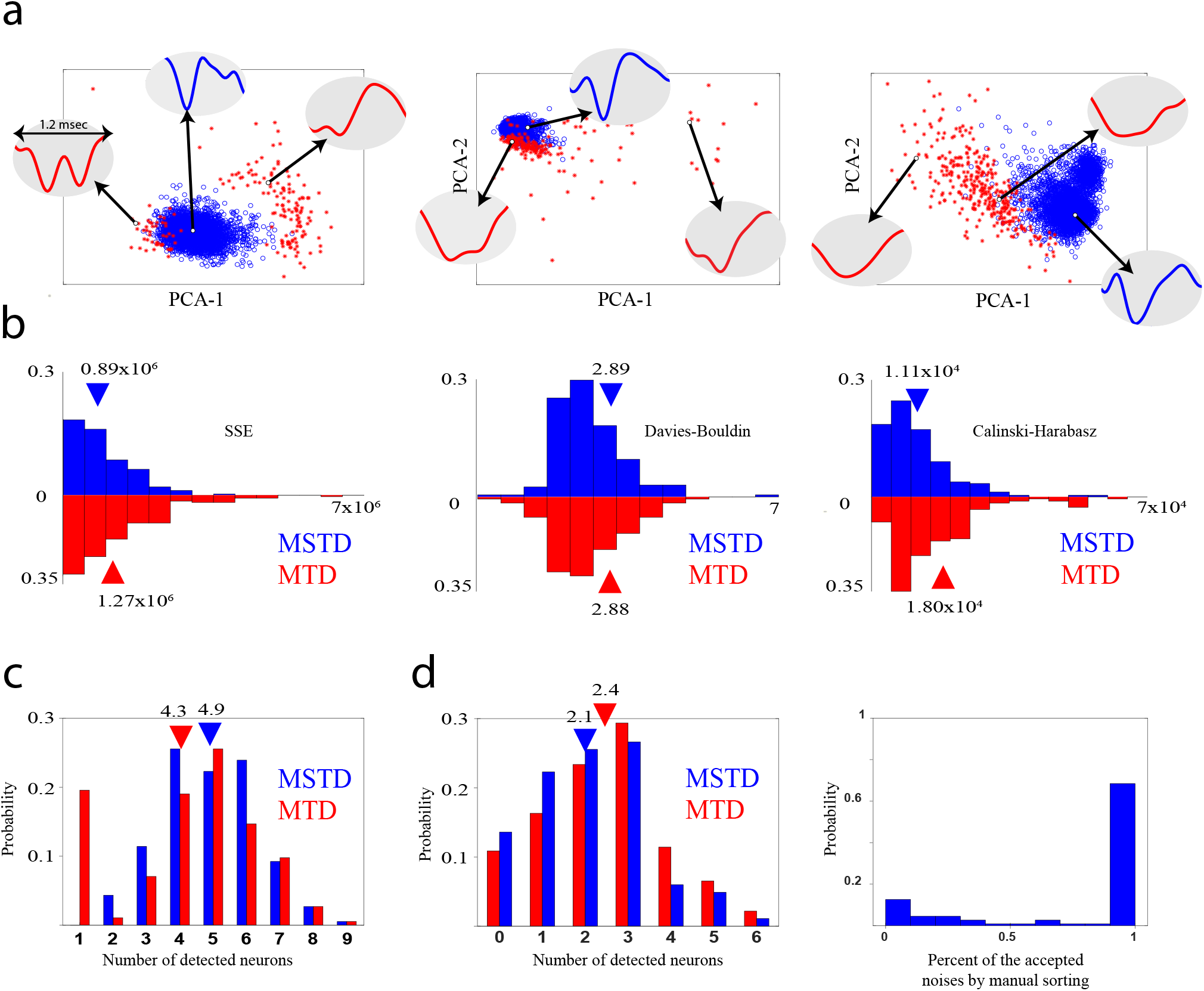
Evaluation of MSTD and MTD using neural recording sessions of the third dataset. (a) Three samples of recording sessions. The waveforms that are detected as noise by statistical filtering are illustrated by red starts and the other waveforms by blue circles. (b) Investigating the quality of clustering using three blind metrics that does not need any ground truth. In the SSE criterion (left), MSTD outperforms MTD, while in Davies-Bouldin criterion (middle) there exists no significant difference; however in terms of Calinski-Harabasz criterion MTD is better. (c) The histogram of the number of clusters determined automatically by each method. (d) Comparison of automatic spike sorting with the manual one. The absolute different between the number of clusters determined automatically by MTD or MSTD and manual sorting (left). The histogram of the percent of the waveforms that are considered as noise both by statistical filtering and manual sorting (right).

Finally to have a better comparison, the resulted clusters of MTD for all sessions have been manually modified by a human operator to achieve a manual sorting outcomes. The available operations are merging, deleting, and resorting the clusters. Also, the results of the statistical filtering, i.e. the waveforms which are detected as noise, are provided for human operator. The operator decides based on the waveform shapes and PCA features of the clusters. Fig. 6(d) shows the absolute difference between the number of clusters determined by the manual sorting and MTD or MSTD methods. The number of clusters in the MSTD method is closer to what is achieved by the manual sorting. We also checked the percent of the samples that are detected as noise by statistical filtering that are removed by the operator in the manual sorting process. Results confirm that in approximately 70% of the sessions, more than 90% of the samples detected as noise by our statistical filtering algorithm are also considered as noise in the manual sorting process.

To sum up the main finding of this section, a comparison between MSTD and MTD methods is summarized in Table 1. The methods are compared regarding the phases where our main findings are bolded, i.e. adaptive detection and clustering, using two datasets: I) synthesized, and II) real dataset with ground truth. Considering the adaptive detection phase, four metrics are used. First, the cluster within distance, reduced by 16%. Second, the number of selected principle components as feature (see equation 4) is decreased by 35%. Finally, the precision and recall are improved by 14% and 17%, respectively, in the real dataset with the ground truth. Considering the clustering phase, the purity and accuracy are improved by 7% and 4% respectively, in the synthesized dataset.

**Table 1.** Summary of the superiority of the MSTD method over MTD. All values in the table are the percent of the improvement (decrease or increase) in the metrics. In the table, D1 and D2 refer to the synthesized and real datasets with ground truth. There exist four metrics that show the effectiveness of our adaptive detection algorithm: I) RCW: reduction in the clusters within distance (see Fig. 5(d)), II) FDR: reduction in the number of principle components selected as feature (see Fig. 5(d)), III) precision: increase in the precision value (see Fig. 5(a), and 3(d)), and IV) recall: increase in recall value (see Fig. 5(a), and 3(d)). In the clustering phase, there exist two metrics: I) purity: increase in resulted clusters’ purity (see Fig. 4(c)), and II) accuracy: increase in accuracy value of the final results (see Fig. 4(c)). The standard error of all values are also presented in the table.

| Dataset | Adaptive Detection |  |  |  | Clustering |  |
| --- | --- | --- | --- | --- | --- | --- |
|  | RCW* (%) | FDR** (%) | precision (%) | recall (%) | purity (%) | accuracy (%) |
| D1 | - | - | $6 \pm 0.1$ | $7 \pm 0.1$ | $7 \pm 4.0$ | $4 \pm 3.1$ |
| D2 | $16 \pm 0.1$ | $35 \pm 0.6$ | $14 \pm 2.8$ | $17 \pm 3.4$ | - | - |

\* Reduction in the cluster within distance
\*\* Feature dimension reduction

## Discussion

In this paper, a novel robust spike sorting algorithm is introduced. In the proposed algorithm, after initial detection in the preprocessing phase, an adaptive detection step is considered to increase the quality of detected spikes. Then, a mixture of skew-t distributions is used for clustering. However, estimating the skewness parameters adds additional computation complexity to the proposed method. The proposed method have three main contributions: i) considering skewness for clusters, new alignment algorithm, iii) statistical filtering for noise removal. It is observed that using the proposed method gives several advantages in both detection and clustering phases. In the detection phase, the correct detection rate is increased while the false alarm rate is decreased. Thus, the MSTD algorithm detects more spikes and less noises. Comparing the automated sorting with a manual sorting, the samples that are detected as noise are also rejected by human operator on roughly 70% of the sessions. Note that noise samples increase the variability of data and could cause automatic sorting algorithms to misclassify them within a neuron or as a new one. Investigating the alignment algorithm shows its effect on the compactness of the spikes of a cluster. As discussed, it reduced the within distance of a cluster, thus makes the clustering easier. In the clustering phase, using artificial dataset, the error shows 40% improvement in high SNRs. Moreover, the purity of clusters increased by roughly 7%. This confirms that the skew-t distributions better fits the feature space of spike waveforms. Another improvement is on the number of clusters which is significantly closer to the real number of clusters according to the simulated and real data with manual sorting.

There exist several challenges for spike sorting algorithms including complex noise^30^, non-Gaussian and skewed cells^31–33^ and time overlapped spikes due to the simultaneous activity of several neurons^34^. We did our best to overcome the first two challenges. While different spike sorting algorithms are developed, yet there exists no universal algorithm that performs well in all situations. Also, the focus of the methods is on the clustering phase. The ISO-SPLIT algorithm is introduced by Chung *et. al.*^35^. Although they introduced a new algorithm for clustering, a simple detection method is used. Besides, the challenge of skewed cells is not investigated at all. The problem of drifting clusters in long recordings is investigated by Shan *et. al.*^18^. They used mixture of drifting t-distribution clustering method to handle this problem. However, authors employ a simple energy operator for spike detection. They stated that their algorithm cannot model cells with skewed distribution. Souza *et. al.* applied Gaussian mixture model (which is a symmetric model that cannot model skewed cells) with variety of features based on PCA and wavelet decomposition^36^ alongside with a threshold crossing event detection. Consequently, their focus is on the feature extraction problem. The challenge of sorting recorded data from high-density multielectrode arrays is studied by Hilgen *et. al.*. In their work, PCA alongside the mean shift algorithm is employed for sorting^37^. Comprehensive sensing is another solution for multielectrode recordings^38^. While PCA is the most common feature extracting method for spike sorting, Caro *et. al.* proposed new feature extraction method based on the shape, phase, and distribution features of each spike^39^. The problem of overlapped spikes could also be handled by applying some post processing on the combination of templates of detected neurons^15, 34, 40^. As can be seen, the focus of the studies has been far from the concerns of this paper, which are robust detection, alignment, and considering skewed cells. By adding an adaptive detection step, and considering skewness in the clustering phase, we add more complexity to the algorithm. However, using parallelism and employing graphics processing unit for implementation could abate the complexity. Another challenge would be the combination of multiple recordings or using different electrodes in a recording, which imposes more complexity. Also spike shape variation may increase false alarm and leads to not well formed clusters. Thus, it is desirable to have an adaptive method dealing with false alarm and not well formed cells.

Our method, sought to fill the gap in the presented spike sorting literature, by targeting and introducing a robust adaptive detection algorithm. In the clustering phase, by applying a mixture of skew-t distribution, the challenge of not well-formed clusters is considered. All in all, our spike sorting algorithm made one step toward robustness against two major challenges: complex noise and non-Gaussian or skewed cells. The implementation of the proposed method is freely available as an open-sourced software. All the mentioned parameters could be modified in the toolbox, which makes it easy to work with any kind of data. The presented toolbox is not merely an offline automatic spike sorting algorithm. It also provides variety of manual functions to improve the quality of the sorting results, such as merging, resorting, removing, manual clustering in the PCA domain, etc. Several visualization functions is also provided to guide the user though the sorting procedure.

## References

1. Sukiban, J. et al. Evaluation of spike sorting algorithms: Application to human subthalamic nucleus recordings and simulations. Neuroscience (2019).

2. Lewicki, M. S. A review of methods for spike sorting: the detection and classification of neural action potentials. Network: Comput. Neural Syst. 9, R53–R78 (1998).

3. Harris, K. D., Henze, D. A., Csicsvari, J., Hirase, H. & Buzsaki, G. Accuracy of tetrode spike separation as determined by simultaneous intracellular and extracellular measurements. J. neurophysiology 84, 401–414 (2000).

4. Kadir, S. N., Goodman, D. F. & Harris, K. D. High-dimensional cluster analysis with the masked em algorithm. Neural computation 26, 2379–2394 (2014).

5. Takekawa, T., Isomura, Y. & Fukai, T. Accurate spike sorting for multi-unit recordings. Eur. J. Neurosci. 31, 263–272 (2010).

6. Takekawa, T., Isomura, Y. & Fukai, T. Spike sorting of heterogeneous neuron types by multimodality-weighted pca and explicit robust variational bayes. Front. neuroinformatics 6, 5 (2012).

7. Hulata, E., Segev, R. & Ben-Jacob, E. A method for spike sorting and detection based on wavelet packets and shannon’s mutual information. J. neuroscience methods 117, 1–12 (2002).

8. Quiroga, R. Q., Nadasdy, Z. & Ben-Shaul, Y. Unsupervised spike detection and sorting with wavelets and superparamagnetic clustering. Neural computation 16, 1661–1687 (2004).

9. Rossant, C. et al. Spike sorting for large, dense electrode arrays. Nat. neuroscience 19, 634 (2016).

10. Wood, F., Black, M. J., Vargas-Irwin, C., Fellows, M. & Donoghue, J. P. On the variability of manual spike sorting. IEEE Transactions on Biomed. Eng. 51, 912–918 (2004).

11. Calabrese, A. & Paninski, L. Kalman filter mixture model for spike sorting of non-stationary data. J. neuroscience methods 196, 159–169 (2011).

12. Carlson, D. E. et al. Multichannel electrophysiological spike sorting via joint dictionary learning and mixture modeling. IEEE Transactions on Biomed. Eng. 61, 41–54 (2013).

13. Franke, F., Natora, M., Boucsein, C., Munk, M. H. & Obermayer, K. An online spike detection and spike classification algorithm capable of instantaneous resolution of overlapping spikes. J. computational neuroscience 29, 127–148 (2010).

14. Shoham, S., Fellows, M. R. & Normann, R. A. Robust, automatic spike sorting using mixtures of multivariate t-distributions. J. neuroscience methods 127, 111–122 (2003).

15. Franke, F., Quiroga, R. Q., Hierlemann, A. & Obermayer, K. Bayes optimal template matching for spike sorting–combining fisher discriminant analysis with optimal filtering. J. computational neuroscience 38, 439–459 (2015).

16. Valencia, D. & Alimohammad, A. An efficient hardware architecture for template matching-based spike sorting. IEEE transactions on biomedical circuits systems 13, 481–492 (2019).

17. Rodriguez, A. & Laio, A. Clustering by fast search and find of density peaks. Science 344, 1492–1496 (2014).

18. Shan, K. Q., Lubenov, E. V. & Siapas, A. G. Model-based spike sorting with a mixture of drifting t-distributions. J. neuroscience methods 288, 82–98 (2017).

19. Quiroga, R. Q. What is the real shape of extracellular spikes? J. neuroscience methods 177, 194–198 (2009).

20. Ison, M. J. et al. Selectivity of pyramidal cells and interneurons in the human medial temporal lobe. J. neurophysiology 106, 1713–1721 (2011).

21. Henze, D. A. et al. Intracellular features predicted by extracellular recordings in the hippocampus in vivo. J. neurophysiology 84, 390–400 (2000).

22. Oppenheim, A. V., Buck, J. R. & Schafer, R. W. Discrete-time signal processing. Vol. 2 (Upper Saddle River, NJ: Prentice Hall, 2001).

23. Cabral, C. R. B., Lachos, V. H. & Prates, M. O. Multivariate mixture modeling using skew-normal independent distributions. Comput. Stat. & Data Analysis 56, 126–142 (2012).

24. Smith, L. S. & Mtetwa, N. A tool for synthesizing spike trains with realistic interference. J. Neurosci. Methods 159, 170–180 (2007).

25. Smith, L. Noisy spike generator, matlab software. Univ. Stirling, Dep. Comput. Sci. Math. (2006).

26. Henze, D. A. et al. Simultaneous intracellular and extracellular recordings from hippocampus region ca1 of anesthetized rats. Available: https://crcns.org/data-sets/hc/hc-1, DOI: 10.6080/K02Z13FP (2009).

27. Henze, D. A. et al. Intracellular features predicted by extracellular recordings in the hippocampus in vivo. J. Neurophysiol. DOI: 10.1152/jn.2000.84.1.390 (2000).

28. Calinński, T. & Harabasz, J. A dendrite method for cluster analysis. Commun. Stat. Methods 3, 1–27 (1974).

29. Davies, D. L. & Bouldin, D. W. A cluster separation measure. IEEE transactions on pattern analysis machine intelligence 224–227 (1979).

30. Einevoll, G. T., Franke, F., Hagen, E., Pouzat, C. & Harris, K. D. Towards reliable spike-train recordings from thousands of neurons with multielectrodes. Curr. opinion neurobiology 22, 11–17 (2012).

31. Quirk, M. C. & Wilson, M. A. Interaction between spike waveform classification and temporal sequence detection. J. neuroscience methods 94, 41–52 (1999).

32. Quirk, M. C., Blum, K. I. & Wilson, M. A. Experience-dependent changes in extracellular spike amplitude may reflect regulation of dendritic action potential back-propagation in rat hippocampal pyramidal cells. J. Neurosci. 21, 240–248 (2001).

33. Harris, K. D., Hirase, H., Leinekugel, X., Henze, D. A. & Buzsáki, G. Temporal interaction between single spikes and complex spike bursts in hippocampal pyramidal cells. Neuron 32, 141–149 (2001).

34. Ekanadham, C., Tranchina, D. & Simoncelli, E. P. A unified framework and method for automatic neural spike identification. J. neuroscience methods 222, 47–55 (2014).

35. Chung, J. E. et al. A fully automated approach to spike sorting. Neuron 95, 1381–1394 (2017).

36. Souza, B. C., Lopes-dos Santos, V., Bacelo, J. & Tort, A. B. Spike sorting with gaussian mixture models. Sci. reports 9, 3627 (2019).

37. Hilgen, G. et al. Unsupervised spike sorting for large-scale, high-density multielectrode arrays. Cell reports 18, 2521–2532 (2017).

38. Xiong, T. et al. An unsupervised compressed sensing algorithm for multi-channel neural recording and spike sorting. IEEE Transactions on Neural Syst. Rehabil. Eng. 26, 1121–1130, DOI: 10.1109/TNSRE.2018.2830354 (2018).

39. Caro-Martín, C. R., Delgado-García, J. M., Gruart, A. & Sánchez-Campusano, R. Spike sorting based on shape, phase, and distribution features, and k-tops clustering with validity and error indices. Sci. reports 8, 17796 (2018).

40. Pachitariu, M., Steinmetz, N., Kadir, S., Carandini, M. & Harris, K. D. Kilosort: realtime spike-sorting for extracellular electrophysiology with hundreds of channels. BioRxiv 061481 (2016).

