## supplementary material for "An Adaptive Detection for Automatic Spike Sorting Based on Mixture of Skew-t distributions"

### Appendix S1: Zero-Phase Filtering

In this method, first the signal is filtered, and then the time reversed version of the filtered signal would be filtered again. Assuming $h$ is the filter response, in the z-transform domain we have:

$$\begin{aligned} &D(z) = R(z)H(z) \\ &R_F(z) = D(z^{-1})H(z) \end{aligned} \tag{S.1}$$

where $R$ is the z-transform of the raw signal $(r)$. When $|z| = 1$ or $z = e^{jw}$, the output reduces to $R_F(e^{jw})|H(e^{jw})|^2$. Thus, the output has zero-phase distortion.

### Appendix S2: Expectation-Maximization for Mixture of Skew-t distribution

In the EM algorithm, it is usual to define a membership variable, $Z_i = (Z_{i1}, ..., Z_{ig})$, where

$$Z_{ij} = \begin{cases} 1, & \text{if } s_i \text{ belongs to the component j} \\ 0, & \text{otherwise} \end{cases} \tag{S.2}$$

The completed log-likelihood function can be written as follows:

$$\begin{aligned} \ell_c(\theta|s,t,u,z) = C - \sum_{i=1}^{N}\sum_{j=1}^{g} Z_{ij} \Big[ &\log(\pi_j) - \frac{1}{2}\log(\gamma_j) - \frac{u_i}{2}(s_i - \mu_j - \Delta_j t_i)^T \gamma_j^{-1}(s_i - \mu_j - \Delta_j t_i) \\ &+ \frac{v}{2}\log(\frac{v}{2}) + (\frac{v}{2} - 1)\log(u_i) - \log(\frac{v}{2}) - \frac{v}{2}u_i \Big] \end{aligned} \tag{S.3}$$

where $C$ is a constant, and

$$\delta = \frac{\lambda_j}{\sqrt{1 + \lambda_j^T \lambda_j}}, \quad \Delta = \Sigma_j^{\frac{1}{2}} \delta_j, \quad \gamma_j = \Sigma_j - \Delta_j \Delta_j^T, \quad t_i \sim N(\mu + \Delta t_i, u_i^{-1}\gamma_j) \tag{S.4}$$

The E-step includes calculating of $Q(\theta|\theta^{(K)}) = E_{\theta^{(K)}}(\ell_c(\theta|s,t,u,z)|s)$, where the superscript $(k)$ indicates the value of the parameters in the $k$'th iteration. The EM algorithm which finds the parameters of a mixture of p-dimensional skew-t distributions can be summarized as follows:

E-Step: First, consider the following auxiliary parameters.

$$
\begin{aligned}
M^2(\theta) &= (1+\Delta^T\gamma^{-1}\Delta)^{-1} \\
m(\theta,s) &= M^2(\theta)\Delta^T\gamma^{-1}(s-\mu) \\
A(\theta,s) &= \lambda^T\Sigma^{-\frac{1}{2}}(s-\mu) \\
\beta(\theta,s) &= \frac{2\Gamma\left(\frac{v+p+2}{2}\right)(v+d_\Sigma(s,\mu))^{-1}}{\Gamma\left(\frac{v+p}{2}\right)} \frac{T\left(\sqrt{\frac{v+p+2}{v-d_\Sigma(s,\mu)}}A(\theta,s)|v+p+2\right)}{T\left(\sqrt{\frac{v-p}{v+d_\Sigma(s,\mu)}}A(\theta,s)|v+p\right)} \\
\tau(\theta,s) &= \frac{1}{\left(\sqrt{\frac{v+p}{v+d_\Sigma(s,\mu)}}A(\theta,s)|v+p\right)} \frac{\Gamma\left(\frac{v+p+1}{2}\right)}{\pi^{\frac{1}{2}}\Gamma\left(\frac{1}{2}\right)} \frac{(v+d_\Sigma(s,\mu))^{(v-p)/2}}{(v+d_\Sigma(s,\mu)+A(\theta,s)^2)^{(v-p+1)/2}} \\
\xi(\theta,s) &= \beta(\theta,s)m(\theta,s)+M(\theta)\tau(\theta,s) \\
\omega(\theta,s) &= \beta(\theta,s)(m(\theta,s))^2+(M(\theta))^2+M(\theta)m(\theta,s)\tau(\theta,s)
\end{aligned} \tag{S.5}
$$

The probability of $i$'th spike belongs to the $j$'th component, i.e. $p_{ij}$, is

$$
p_{ij} = \frac{\pi_j ST_p(s_i|\theta_j)}{\sum_{l=1}^{g}\pi_l ST_p(s_i|\theta_l)} \tag{S.6}
$$

Given $\theta=\theta^{(k)}$, first we calculate $p_{ij}^{(k)}$, then,

$$
\beta_{ij} = p_{ij}^{(k)}\beta(\theta_j^{(k)},s_i) \quad \xi_{ij} = p_{ij}^{(k)}\xi(\theta_j^{(k)},s_i) \quad \omega_{ij} = p_{ij}^{(k)}\omega(\theta_j^{(k)},s_i) \tag{S.7}
$$

M-Step: In this step, by maximizing $Q(\theta|\theta^{(k)})$, the update rules for the parameters would be obtained as follows:

$$
\begin{aligned}
\pi_j^{(k+1)} &= n^{-1}\sum_{i=1}^{N} p_{ij}^{(k)} \\
\mu_j^{(k+1)} &= \sum_{i=1}^{N}(\beta_{ij}^{(k)}s_i-\xi_{ij}^{(k)}\Delta_j^{(k)})/\sum_{i=1}^{N}\beta_{ij}^{(k)} \\
\Delta_j^{(k+1)} &= \left[\sum_{i=1}^{N}\xi_{ij}^{(k)}(s_i-\mu_j^{(k+1)})\right]/\sum_{i=1}^{N}\omega_{ij}^{(k)} \\
\gamma_j^{(k+1)} &= \left(\sum_{i=1}^{N}p_{ij}\right)^{-1}\sum_{i=1}^{N}\left(\beta_{ij}^{(k)}(s_i-\mu_j^{(k+1)})(s_i-\mu_j^{(k+1)})^T\right. \\
&\quad \left.-\left[(s_i-\mu_j^{(k+1)})(\Delta_j^{(k+1)})^T+(\Delta_j^{(k+1)})(s_i-\mu_j^{(k+1)})^T\right]\xi_{ij}^{(k)}+\Delta_j^{(k+1)}(\Delta_j^{(k+1)})^T\omega_{ij}^{(k)}\right) \\
v^{(k+1)} &= \arg\max_{v}\sum_{i=1}^{N}\log\left(\sum_{j=1}^{g}\pi_j ST_p(s_i|\mu_j^{(k+1)},\Sigma_j^{(k+1)},\lambda_j^{(k+1)},v^{(k)})\right)
\end{aligned} \tag{S.8}
$$

The update procedure is based on $\Delta$, and $\gamma$ instead of $\lambda$ and $\Sigma$. These parameters could be recovered as follows:

$$
\lambda_j = \frac{(\gamma_j+\Delta_j\Delta_j^T)^{-\frac{1}{2}}\Delta_j}{\left[1-\Delta_j^T(\gamma_j+\Delta_j\Delta_j^T)^{-1}\Delta_j\right]^{\frac{1}{2}}} \qquad \Sigma_j = \gamma_j+\Delta_j\Delta_j^T \tag{S.9}
$$
